# Recurrent plant-pathogen Enterobacterales offer complementary digestive functions in a polyphagous insect pest, *Empaosca fabae*

**DOI:** 10.64898/2026.08.08.743697

**Authors:** Joshua Molligan, Thierry Pellegrinetti, Elisa Ines Fantino, Edel Pérez-López

## Abstract

Nutritional homeostasis in many leafhoppers (Cicadellidae) is largely attributed to ancient obligate symbionts, yet the facultative bacteria these insects carry and if whether they contribute to digestion, remains poorly understood. This question is especially relevant in mesophyll cell-rupture feeders of the subfamily Typhlocybinae, which are reported to lack classical obligate associations. The potato leafhopper, *Empoasca fabae*, is a polyphagous, migratory Typhlocybine that feeds on more than 200 plant species. Metagenomic analysis of field-collected *E. fabae* recovered four complete metagenome-assembled genomes corresponding to the opportunistic plant-pathogenic Enterobacterales *Enterobacter mori, Kosakonia cowanii, Pantoea agglomerans*, and *Pantoea ananatis*, each highly similar to its type strain. Species-specific PCR across a five-year window showed that *E. mori* and *K. cowanii* were detected in every field sample and persistent in an inbred colony, demonstrating likely recurrent and maintained associations, whereas the two *Pantoea* species were detected intermittently. All four genomes encoded broad carbohydrate-processing repertoires, including sucrose phosphotransferase systems, glycolysis, and aromatic amino acid biosynthesis, suggesting a capacity to synthesize aromatic amino acids–essential for the host. Among 614 glycoside hydrolases, two putatively secreted GH5-25 cellulases were further examined, with recombinant *K. cowanii* KcGH5-1 hydrolyzing carboxymethyl cellulose at acidic pH, signifying a functional bacterial endoglucanase. These results identify recurrent plant-pathogenic Enterobacterales as carriers of complementary digestive functions, and as candidate contributors to the exceptional dietary breadth of a major migratory agricultural pest.

---

Leafhoppers (Cicadellidae) are among some of the most diverse plant-feeding insects, with their nutrition typically supported by ancient obligate symbionts [1, 2]. However, the facultative bacteria they carry, and whether these contribute to digestion, remain poorly understood [3]. Facultative associates are of particular interest in mesophyll feeders of the subfamily Typhlocybinae, which have been reported to lack obligate partners entirely [4]. The potato leafhopper, *Empoasca fabae* (Harris, 1841), is an unusually polyphagous Typhlocybine that feeds on more than 200 plant species and migrates annually across much of eastern North America [5–7]. Whether its microbiome carries carbohydrate-processing capacity that could complement this diet in the absence of obligate members remains unknown.

To characterize the *E. fabae* microbiome, we collected eighteen pooled field samples (ten individuals each) in mid-August 2023 from a single strawberry farm in southern Québec, Canada, the northern extent of *E. fabae*’s migratory range [8]. We then performed shotgun metagenomic sequencing and recovered metagenome-assembled genomes (MAGs) for the dominant taxa. This resolved four complete MAGs (100% completeness, <0.4% contamination, 4.68–4.82 Mb) corresponding to *Enterobacter mori, Kosakonia cowanii, Pantoea agglomerans*, and *Pantoea ananatis*, each with >97.8% average nucleotide identity (ANI) to its type strain (**Fig. 1A, Data S1**). All four have been described independently as opportunistic plant pathogens of crops including soybean, kiwifruit, walnut, and rice, none of which, however, have been reported as pathogens in the fields surrounding this collection [9–12]. Their persistent recovery as high-quality genomes from the *E. fabae* metagenome suggested a stable, potentially functional association (**Data S2**).

**Figure 1.**
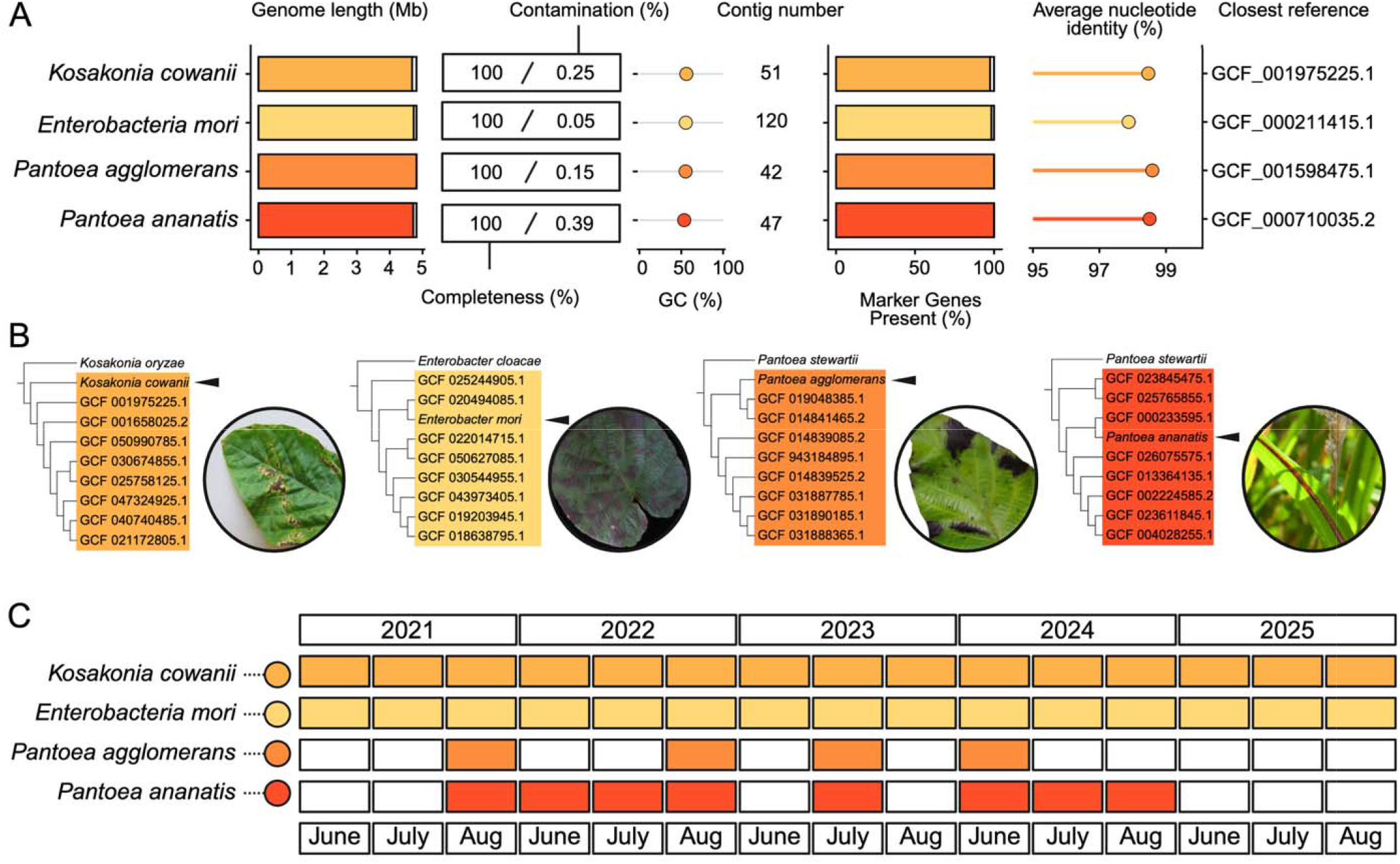
Recurrent Enterobacterales part of *Empoasca fabae* microbiome. (**A**) Quality metrics and taxonomic placement of the four facultative Enterobacterales MAGs recovered from the *E. fabae* metagenome: genome length, completeness, contamination, GC content, contig number, marker genes present, and average nucleotide identity (ANI) to the closest reference genome. (**B**) Phylogenetic placement of the four Enterobacterales MAGs compared to representative type strains of each genus. Each species is a documented plant-opportunistic pathogen (representative disease-symptom photographs shown). (C) Presence/absence of the four Enterobacterales across *E. fabae* samples screened by species-specific PCR (June-August, 2021-2025). Field samples (2021-2024) were collected in the Montérégie region, Québec, Canada. Samples from 2025 represent the inbred laboratory colony tested. Disease-symptom photographs are reproduced under Creative Commons licenses (CC0 1.0 and CC BY 4.0; see Supplementary Information).

To test this persistence, we screened field-collected and colony-reared *E. fabae* for all four taxa using species-specific diagnostic PCR across a broader five-year window (2021–2025) from the same collection site (**Fig. 1B, Data S3**). *E. mori* and *K. cowanii* were detected in every field sample and persisted in an inbred laboratory colony started from insects captured at the original site. Colony samples were tested regularly throughout the first year of colony establishment, spanning approximately 10 insect generations, after which they consistently tested negative. This suggests a recurrent, maintained association that appears to degenerate without external environmental acquisition. *Pantoea agglomerans* and *P. ananatis* were detected intermittently across field samples, but never persisted in the colony, consistent with environmentally acquired associates. We suggest that together, all presented bacteria represent associates acquired environmentally, before or after migration, with yet unknown factors influencing retention. To our knowledge, stable, multi-year retention of facultative Enterobacterales has not previously been reported in Cicadellidae.

As plant-associated bacteria, we then asked whether these taxa encoded functions relevant to a phytophagous diet. Reconstruction of KEGG metabolic modules across the four MAGs showed broad and largely shared enrichment in starch and sucrose metabolism, glycolysis/gluconeogenesis, and aromatic amino acid biosynthesis (phenylalanine, tyrosine, tryptophan) (**Fig. 2A**). Each genome encoded sucrose-specific phosphotransferase systems that permit glucose-6-phosphate to commence central carbon metabolism, together with secretion machinery (Sec, Tat) and complete shikimate-to-chorismate pathways (**Fig. 2B**). Together, these findings suggest that each associate bacteria can catabolize plant-derived sugars and synthesize aromatic amino acids. These traits are suspected of plant-associated bacteria, however given their recurrent presence we hypothesize they may also serve to compensate for limitations in insect amino acid metabolism.

**Figure 2.**
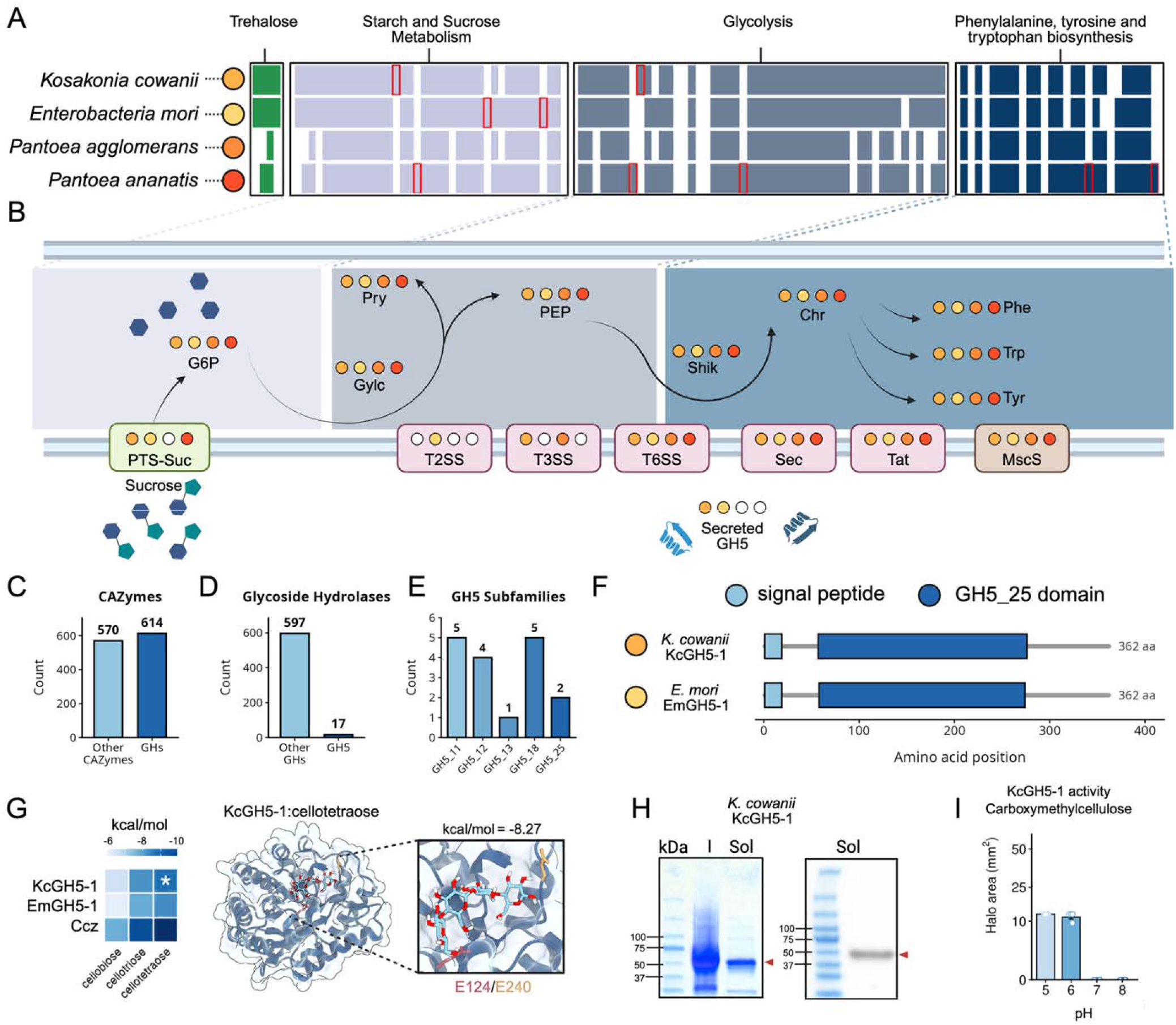
Carbohydrate-processing capacity and a functional secreted cellulase in *E. fabae*-associated Enterobacterales. (**A**) KEGG metabolic pathway completeness across the four Enterobacterales MAGs for four categories: trehalose metabolism, starch and sucrose metabolism, glycolysis/gluconeogenesis, and aromatic amino acid biosynthesis (phenylalanine, tyrosine, tryptophan). Steps are shown as present (filled) or absent (white). Red boxes highlight unique enzymatic steps to a single MAG. (**B**) Central carbon metabolism and transport organization, showing sucrose processing, glycolysis, and the phosphoenolpyruvate (PEP)-to-shikimate branch, with phosphotransferase (PTS) transporters and secretion systems (Sec, Tat, MscS). Steps are colored the corresponding color of a MAG if considered fully complete by manual inspection. The color used for each MAG matches the color scheme used in panel **A** and in **Figure 1**. (**C**) Number of other CAZyme families versus glycoside hydrolases (GHs). (**D**) Number of other GHs compared to the number of GH5s. (**E**) Number of GH5 by subfamily (GH5-11, GH5-12, GH5-13, GH5-18, and GH5-25) (**F**) Domain architecture of the two secreted GH5-25 cellulases, EmGH5-1 (*E. mori*) and KcGH5-1 (*K. cowanii*), each with a predicted N-terminal signal peptide and a catalytic domain. (**G**) Molecular docking of KcGH5-1, EmGH5-1, and the control CcsZ (*C. difficile*) to representative β-1,4-glucan substrates. The strongest-scoring KcGH5-1 pose is shown with a cellotetraose ligand in the catalytic cleft and catalytic residues E124/E240 highlighted. (**H**) Heterologous expression and Ni-NTA purification of soluble His-tagged KcGH5-1 [pET-28a(+), *E. coli*; I, induced; Sol, soluble fraction; red arrowhead]. (**I**) Halo degradation areas of purified KcGH5-1 on carboxymethylcellulose (CMC) agar at pH 5 to 8.

As a mesophyll-feeder exposed to a diversity of plant structural polysaccharides, *E. fabae* may benefit from bacterial cell wall-degrading capacity, we therefore further profiled carbohydrate-active enzymes (CAZymes) across the four associate bacteria genomes. We identified 1,184 CAZymes, of which 614 (52%) were glycoside hydrolases (GHs) (**Fig. 2C**). Among these were 17 GH5 enzymes spanning five subfamilies, including two GH5-25 cellulases carrying N-terminal signal peptides, KcGH5-1 (*K. cowanii*, PIEDBF_04288) and EmGH5-1 (*E. mori*, IMICMD_01186), indicating secreted, extracellular endo-β-1,4-glucanase activity with *in silico* substrate docking support (**Fig. 2D-G, Fig. S1, S2, Data S4**). To test whether these predicted cellulases are functional, we expressed both enzymes in *Escherichia coli*, with the characterized GH5-25 CcsZ (*Clostridioides difficile*) as a control, and assayed activity on carboxymethyl cellulose (CMC) across pH 5–8 (**Fig. 2H, Fig. S3, Data S5**). Recombinant KcGH5-1 hydrolyzed CMC, producing clear degradation halos at pH 5 and 6 with no detectable activity above pH 6, indicating an acidic-optimum endo-β-1,4-glucanase (**Fig. 2I**). No activity was detected for EmGH5-1 under any condition, despite confirmed soluble expression (**Fig. S4A**). Nonetheless, this demonstrates a recombinant enzymatically active GH5-25 on cellulose from a recurrent *E. fabae*-associated Enterobacterales, extending a suspected facultative nutritional role beyond the presented predictive metabolic reconstructions.

Despite repeated attempts we were unable to isolate *E. mori* or *K. cowanii* in axenic culture from surface-sterilized insects. Therefore, the presented genomic and enzymatic evidence relies on metagenomic and recombinant data rather than cultured strains. Additionally, KcGH5-1 activity was observed *in vitro* under acidic conditions, and its expression and deployment *in vivo* remain to be established as the enzyme could be conditionally active in contexts not captured here, similar to absence of activity in EmGH5-1.

Our previous genome-resolved survey of 171 leafhopper species revealed a recurrent modular microbiome architecture comprising conserved obligate symbionts, a flexible layer of secondary symbionts, and a dynamic pool of environmentally acquired bacteria [13]. Here, we begin to resolve the functional significance of this dynamic layer in *E. fabae*. Recurrently associated Enterobacterales encode complementary repertoires for sugar catabolism and polysaccharide degradation, and the biochemical validation of KcGH5-1 demonstrates that at least one of their predicted glycoside hydrolases is an active cellulase. This finding moves the accessory microbiome beyond genomic prediction and identifies a testable mechanism through which environmentally acquired bacteria may expand the range of plant carbohydrates accessible to *E. fabae*. Such metabolic flexibility may help support the exceptional polyphagy of this migratory pest, providing a direct link between microbiome composition, digestive capacity, and dietary breadth.

## ACKNOWLEDGEMENTS

We are thankful to Abrãao Almeada Santos and Jordanne Jacques who helped collect insects for sequencing. Figures were assembled in BioRender.com.

## AUTHOR CONTRIBUTIONS

Conceptualization, J.M. and E.P.-L.; Methodology, J.M., T.P., E.I.F.; Investigation, J.M. and E.P.-L.; Data curation and formal analysis, J.M.; Visualization, J.M.; Resources and funding acquisition, E.P.-L.; Project administration and supervision, E.P.-L.; Writing original draft, J.M., T.P., and E.P.-L; All authors contributed to review & editing of this manuscript.

## DATA AVAILABILITY

The sequencing datasets generated for this study are publicly available through NCBI under the following BioProject accessions: PRJNA1160200/PRJNA1442128 (*Empoasca fabae* mitogenome/metagenome sequencing and assembly). All presented MAGs and associated annotaions are available at Zenodo (https://doi.org/10.5281/zenodo.21828961)

