## Supplementary files for "Recurrent plant-pathogen Enterobacterales offer complementary digestive functions in a polyphagous insect pest, *Empaosca fabae*"

### SUPPLEMENTARY METHODS

#### Field-Collected *E. fabae* Adults.

Adult *E. fabae* were collected from an agricultural site in southern Québec, Canada from the Montérégie region. From 2021-2024, collections from a single agricultural field in Montérégie was performed using yellow sticky traps, and specimens were preserved in 70% ethanol at 4°C until processing. This collection was part of previous studies [1, 2]. The DNA obtained from these specimens was used for metagenomic sequencing and PCR screenings. Specimens from all field collection events have been deposited at the Canadian National Collection of Insects, Arachnids, and Nematodes, under the voucher numbers CNC2098398-2098407, with Dr. Joel Kits as the person responsible of the Hemiptera division. Methods describing COI-based species verification of all metagenomic samples are provided below. Morphological identification of *E. fabae* was performed by co-authors Molligan J. and Plante N. as described in previous studies [1, 2].

#### *E. fabae* Colony Establishment and Diet Assay Experimental Design.

A colony of *E. fabae* was maintained under controlled conditions at Université Laval reared on alfalfa plants beginning in July of 2024. Insects were maintained at 21°C during the day and 18°C at night on an 18:6 h light:dark cycle. Host plants were fertilized weekly and replaced every three to four weeks.

#### Additional COI-Based Species Verification.

Paired-end reads from each of the 18 pool-sequenced libraries were quality-trimmed with fastp v1.0. Trimmed reads were mapped to a BOLD Cicadellidae reference database (all Canadian and US Cicadellidae records = 40,379 barcodes) supplemented with the *E. fabae* mitogenome COI (NCBI PQ351619.1) using Bowtie2 v2.5.4. COI-aligned reads were extracted with SAMtools v1.17 and assembled per sample with SPAdes v3.15.5. Resulting contigs were queried by BLASTN against the same reference. The top matches of each contig were to the *E. fabae* mitogenome sequence at ≥99.94% identity.

**Symbiont Metagenome Reconstruction.**

The genomic DNA of eighteen field collected *E. fabae* samples consisting of 10 pooled individuals was extracted as previously described [1]. The DNA was then sequenced using Illumina paired-end sequencing (Genome Québec, CA). Reads were adapter and quality trimmed with BBTools v36.92 [3] and aligned to the *E. fabae* genome reference genome (GCA_057929975.1), using bwa-mem2 v2.3. Unaligned reads were retained and considered as the microbial fraction. Insect-filtered reads were assembled using metaSPAdes v3.9.0 [4]. Contigs were binned using MaxBin2 v2.2.7 [5], MetaBAT2 v2.18 [6], and CONCOCT v1.1 [7] then consolidated with DAS Tool v1.1.7 [8]. Metagenome-assembled genomes (MAGs) from all samples were dereplicated using dRep v3.4.2 [9]. Genome completeness and contamination were evaluated with CheckM2 1.1.0 [10], and taxonomic classification was performed using GTDB-Tk 2.5.2 [11] with GTDB-Tk release data r226 [12]. Relative abundance and coverage were estimated by mapping insect-depleted reads back to all dereplicated MAGs with Bowtie2 v2.5.4 [13]. Mean depth was calculated using SAMtools v1.17. Further taxonomic refinement and phylogenetic placement of MAGs within plant associated genera was done by comparing conserved orthologs against ten RefSeq genomes. Orthologous genes were identified using OrthoFinder v3.1.0, then aligned with MAFFT v7.533 and trimmed using trimAl v1.4. A phylogenetic tree was then constructed with IQ-TREE2 v3.0.1 with the best-fit substitution model selected by ModelFinder.

**Metabolic Reconstruction of Symbionts.**

All MAGs were annotated using Bakta v1.11.4 [14]. Amino acid sequences were then used to assign KEGG orthologs (KOs) (107, 108) using GhostKOALA [15]. KO profiles were parsed to evaluate starch and sucrose metabolism, glycolysis, gluconeogenesis and essential amino acid biosynthesis modules. Pathway completeness was manually evaluated stepwise according to representative KEGG modules, considering only absolute presences for module completion.

**Recurrent Enterobacterales Screening.**

Field-collected and colony reared *E. fabae* adults were screened for four recurrent Enterobacterales associates (*Enterobacter mori*, *Kosakonia* cowanii, *Pantoea ananatis*, and *Pantoea agglomerans*) using species-specific PCR assays. Primer sets were selected from published diagnostic protocols and initially aligned to respective MAGs to confirm sequence identity. For *E. mori*, we used the *rpoB*-based assay with primers pair Em-rpoBF/Em-rpoBR [16]. For *P. ananatis*, we used the *gyrB*-targeting primers pair PANAN_gyrB_fwd/PANAN_gyrB_rev [17]. For *P. agglomerans*, we used the infB-targeting primers pair PANAG_infB_fwd/PANAG_infB_rev [17]. For *K. cowanii*, we used species-specific primers pair KC_fwd/KC_rev [18]. Each PCR assay was performed using *E. fabae* gDNA as template, following the recommended conditions and protocols. Amplicon sizes were confirmed through electrophoresis on agarose gel, and products of the expected size were sequenced to confirm amplicon identity (CHUL, Québec, CA). Insects were considered positive for a given symbiont when the expected-size amplicon was obtained.

**Microbial CAZyme and GH Annotation.**

CAZymes were annotated across all MAGs using dbCAN3 with the dbCAN database [19]. GH families were extracted from dbCAN results to capture both family-level and subfamily-level classifications. Gene-level counts were then aggregated across MAGs and compiled into family-wide and subfamily-wide counts. All GH5 enzymes were then screened for signal peptides using SignalP v6.0 [20]. Enzymes containing N-terminal signal peptides were prioritized for functional characterization. Domain architectures of secreted GH candidates were further validated using InterProScan v5.0 and compared against the UniProt database [21, 22].

**GH Structural Modeling and Molecular Docking.**

Three-dimensional structures of GH5-25 enzymes were predicted using AlphaFold3 [23]. Model quality was assessed using per-residue predicted local distance difference test (pLDDT) scores and predicted aligned error (PAE) matrices. Structural analysis was performed in UCSF ChimeraX v1.11.1 [24]. Structural superimpositions between domain architectures were computed. Active-site geometry and catalytic residue positioning were examined and compared between domains.

Oligosaccharide substrates representing major GH5-25 relevant classes were constructed using GLYCAM-Web [25], including β-1,4-glucans (cellobiose, cellotriose, cellotetraose), β-1,4-mannans (mannotriose), galactomannans (galacto-mannotriose), and xyloglucan oligosaccharides. Ligand geometries were energy-minimized and compiled into an SDF library using OpenBabel v3.1.1 [26]. Docking simulations were performed using GNINA [27], a CNN-enhanced AutoDock Vina derivative. Receptor structures were prepared by adding polar hydrogens, and the docking search space was defined using GNINA's autobox procedure centered on the catalytic cleft. Docking employed exhaustiveness of 32 with a fixed random seed for reproducibility. Binding pose analysis was visually inspected in ChimeraX v1.11.1.

**GHs Cloning and Heterologous Expression.**

Coding sequences of microbial GH5-25 enzymes (*C. difficile* CcsZ, *K. cowanii* KcGH5-1, and *E. mori* EmGH5-1), without signal peptides, were codon-optimized for *Escherichia coli* expression, synthesized and cloned into pET-28a(+) vectors with N-terminal 6xHis tags (Twist Bioscience, San Francisco, California, USA). Expression vectors were transformed into *E. coli* LEMO21(DE3) cells (New England Biolabs, Ipswich, Massachusetts, USA) by heat shock transformation. Transformants were selected on LB agar containing kanamycin (50 µg/mL).

Single colonies were inoculated into LB medium supplemented with kanamycin (50 µg/mL) and chloramphenicol (34 µg/mL) and grown at 37°C with shaking (220 rpm) to mid-log phase ($OD_{600}$= 0.5-0.8). Protein expression was induced with 0.5 mM IPTG, followed by incubation at 16°C for 18 h. Cells were harvested by centrifugation at 4°C, resuspended in lysis buffer (50 mM $NaH_2PO_4$,

300 mM NaCl, 10 mM imidazole, pH 8.0), and disrupted by sonication (6 cycles of 30 s on/30 s off at 35% amplitude using Fisherbrand Model 505 Sonic Dismembrator; Thermo Fisher Scientific, Waltham, Massachusetts, USA). The soluble fraction was recovered by centrifugation (12,000 x g, 30 min, 4°C) and purified using Ni-NTA agarose chromatography (QIAGEN, Hilden, Germany) according to manufacturer's protocols. Bound proteins were eluted with increasing imidazole concentrations (50-250 mM). Proteins were quantified using Quick Start Bradford Protein Assay (Bio-Rad, Hercules, California, USA) and stored at 4°C.

Protein purity was assessed by SDS-PAGE using 4-15% Mini-PROTEAN TGX precast gels (Bio-Rad) stained with Bio-Safe Coomassie (Bio-Rad). For Western blotting, proteins were transferred to 0.2 µm PVDF membranes (70 V, 400 mA, 2 h), blocked with 5% milk (1 h, RT), and washed three times with TBST (Tris-buffered saline, 0.1% Tween 20). Anti-$His_6$-HRP conjugate antibody (Roche, Basel, Switzerland; Ref 11965085001, lot 47120100) was used at 1:500 dilution in 3% BSA and incubated for 1 h at RT. After three TBST washes, proteins were detected using Clarity Max Western ECL substrate (Bio-Rad). Chemiluminescence was visualized using an Azure c300 imaging system with AzureSpot software (Azure Biosystems, Dublin, California, USA). Ponceau S staining (5% glacial acetic acid, 0.1% Ponceau Red dye) served as a loading control.

**Polysaccharide Activity Assays.**

Endo-β-1,4-glucanase and endo-β-1,4-mannanase activities were assessed using agar plate diffusion assays. Substrate plates were prepared by dissolving either carboxymethylcellulose sodium salt (CMC; 0.2% w/v, Thermo Fisher Scientific) or locust bean gum (LBG; 0.2% w/v, Thermo Fisher Scientific) in agar (1.5% w/v) adjusted to pH 5, 6, 7, or 8 using appropriate buffer (50 mM sodium citrate for pH 5 and 6; 50 mM sodium phosphate for pH 7 and 8). Equal volume droplets of 100 mM of purified protein or buffer-only controls were placed on agar surfaces. Plates were incubated at 28°C for 24 h to test for hydrolysis. Following incubation, plates were then stained with 0.1% (w/v) Congo red for 15 min and destained with 1 M NaCl to reveal degradation zones. Plates were further neutralized with 1% (v/v) HCl to enhance halo contrast. Cleared halos surrounding each spot were measured and halo areas ($mm^2$) were quantified.

**Plant pathogen symptomology photos in Figure 1. credited to:**

*Pantoea agglomerans* walnut pathogenesis photo:

J. Xiao, et al., Complete genome sequence of *Pantoea agglomerans* CHTF15, a walnut pathogen. *Mol. Plant Microbe Interact.* 36, 134–137 (2023).

*Pantoea ananatis* rice infection photo:

R. Pedrozo, et al., New threats to rice production: Emerging pathogens and their impact. In *Rice Production Strategies* (IntechOpen, 2025).

*Kosakonia cowanii* soybean pathogenesis photo:

K. Krawczyk, N. Borodynko-Filas, *Kosakonia cowanii* as the new bacterial pathogen affecting soybean (*Glycine max*Willd.). *Eur. J. Plant Pathol.* 157, 173–183 (2020).

*Enterobacter mori* kiwifruit pathogenesis photo:

M. Zhang, et al., Whole genome sequencing of *Enterobacter mori*, an emerging pathogen of kiwifruit and the potential genetic adaptation to pathogenic lifestyle. *AMB Express* 11, 129 (2021).

### SUPPLEMENTARY FIGURES

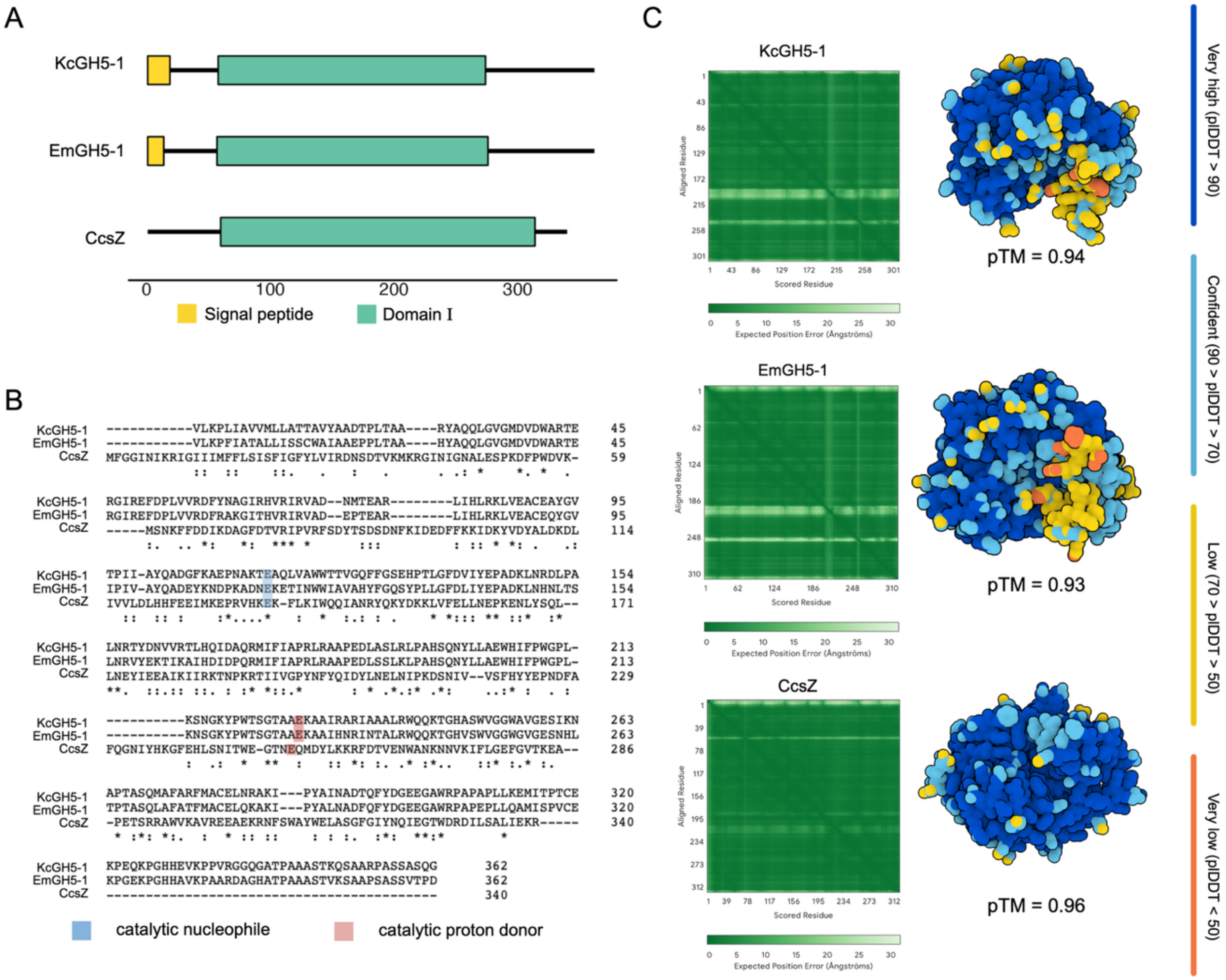


**Figure S1.** Domain architecture, sequence alignment, and predicted structures of GH5-25 enzymes (KcGH5-1, EmGH5-1, CcsZ). (***A***) Schematic domain architecture of three GH5-25 enzymes from leafhopper-associated bacteria with signal peptides: *K. cowanii* KcGH5-1, *E. mori* EmGH5-1, and *C. difficile* CcsZ. Protein lengths are drawn to scale with signal peptides shown in yellow and GH5-25 domains in teal. (***B***) Multiple sequence alignment of the three GH5-25 enzymes. Conserved catalytic nucleophile (blue) and proton donor (pink) residues are highlighted. Consensus symbols indicate fully conserved (*), strongly similar (:), and weakly similar (.) positions. (***C***) AlphaFold3-predicted structures of the full-length modular proteins. Left: predicted aligned error (PAE) matrices, where light green indicates low expected positional error between residue pairs and dark green coloring indicates high confidence in relative positioning. Right: surface representations colored by per-residue pLDDT confidence. Predicted template modeling (pTM) scores are indicated for each structure.

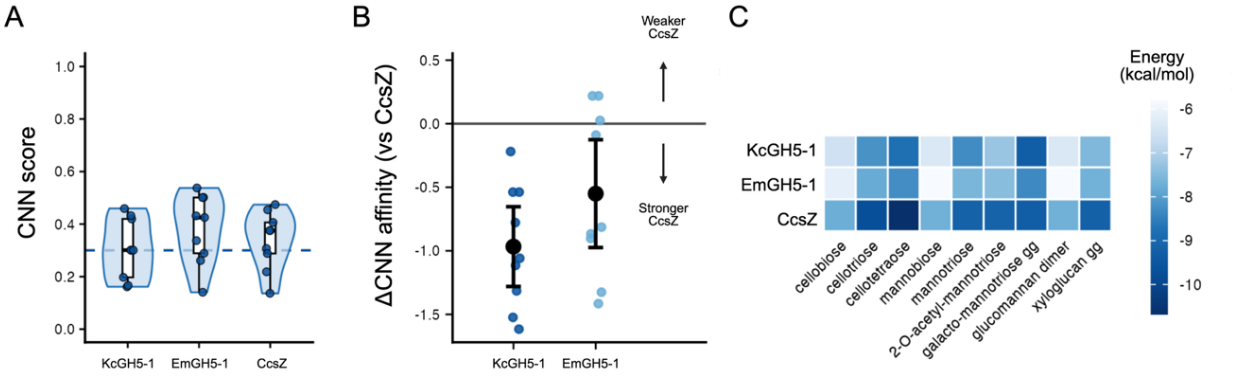


**Figure S2.** *In silico* substrate interaction predictions of KcGH5-1, EmGH5-1, and CcsZ. (***A***) CNNscore distributions for KcGH5-1, EmGH5-1, and CcsZ across all docked ligands. Violins show the full score distribution and points show individual docking poses. Dashed line indicates a CNNscore threshold of 0.3. (***B***) Differential CNNaffinity of KcGH5-1 and EmGH5-1 relative to CcsZ across ligands. Points represent individual ligands and black circles demonstrate the mean with error bars. Negative values indicate stronger predicted binding by CcsZ. (***C***) Heatmap of binding energies (kcal/mol) for all three proteins across ligands. Darker blue indicates more negative (stronger) predicted binding energy.

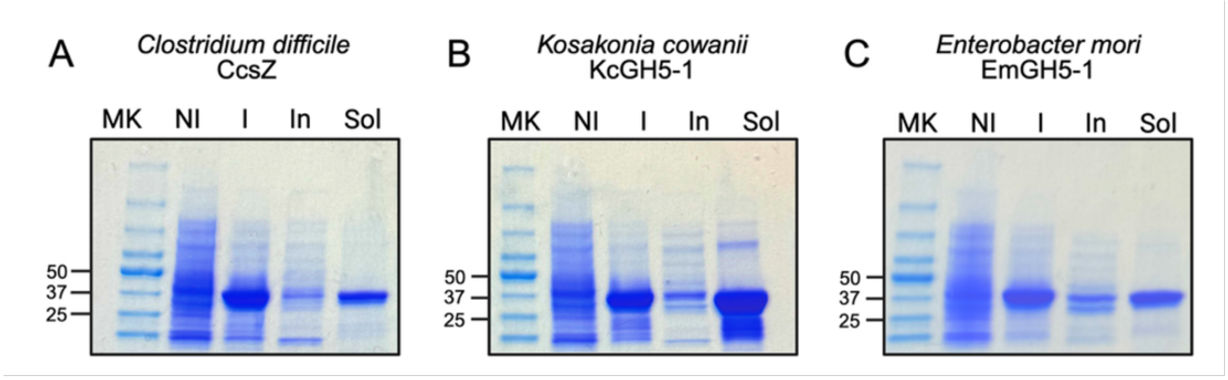


**Figure S3.** SDS-PAGE of recombinant GH5-25. (*A-C*) SDS-PAGE of cell lysate fractions from *E. coli* Lemo21(DE3) expressing recombinant GH5 enzymes. Lanes: MK, molecular weight marker (kDa); NI, non-induced whole-cell lysate; I, IPTG-induced whole-cell lysate; In, insoluble pellet fraction; Sol, soluble supernatant fraction. Expected molecular weights are as follows: CcsZ, 44 kDa; KcGH5-1, 42 kDa; EmGH5-1, 41.3 kDa.

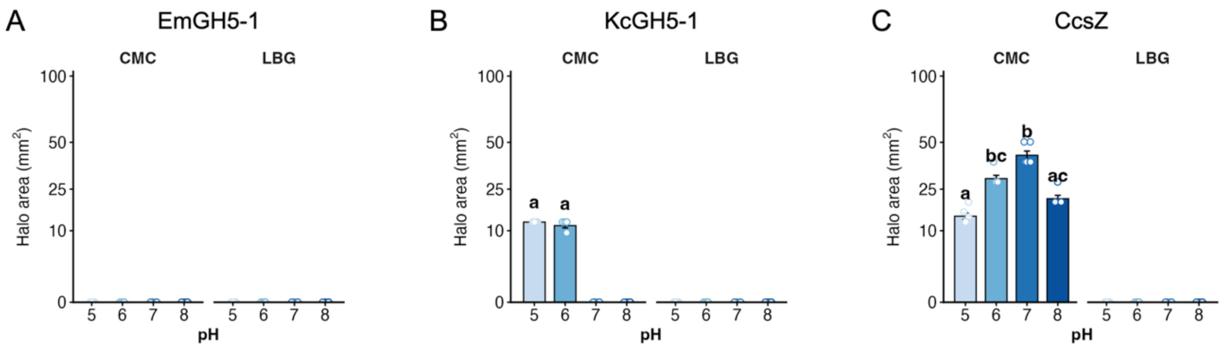


**Figure S4.** KcGH5-1, EmGH5-1, and CcsZ halo activity towards representative substrates. (***A-C***) Halo area ($mm^2$) for EmGH5-1 (***A***), KcGH5-1 (***B***), and CcsZ (***C***) on carboxymethyl cellulose (CMC) and locust bean gum (LBG) media at pH 5, 6, 7, and 8. Bars show mean ± SE and points show individual replicates (*n* = 6). Letters indicate statistically significant differences between pH conditions within each media (Kruskal-Wallis with Dunn's post-hoc test, BH-adjusted, groups sharing a letter are not significantly different). Y-axis is square-root transformed and labels reflect original $mm^2$ values.
